# Tunable Cellular Localization and Extensive Cytoskeleton-Interplay of Reflectins

**DOI:** 10.1101/2021.08.23.457345

**Authors:** Junyi Song, Chuanyang Liu, Baoshan Li, Liangcheng Liu, Ling Zeng, Zonghuang Ye, Ting Mao, Wenjian Wu, Biru Hu

## Abstract

Reflectin proteins are nature copolymers consist of repeated canonical domains, locating in a biophotonic system called Bragg lamellae and manipulating the dynamic structural coloration of iridocytes. Their biological functions are intriguing, but the underlying mechanism is not fully understood. In this study, reflectin A1, A2, B1 and C were found to present distinguished cyto-/nucleoplasmic localization preferences. Comparable intracellular localization can be reproduced by truncated reflectin variants, suggesting a conceivable evolutionary order among reflectin proteins. Secondly, the size-dependent access of reflectin variants into nucleus demonstrate a potential model of how reflectins get into Bragg lamellae. Thirdly, RfA1 was found to extensively interact with cytoskeleton, including its binding to actin and microtubule minus-end-directed movement. This implies cytoskeleton system plays fundamental role during the organization and transportation of reflectin proteins. Findings presented here recommend reflectins as programmable biomaterials which can be used to decipher their evolution processes, to delineate their biological mechanism, and to achieve tunable intracellular targeting as editable tags.

## Introduction

Cephalopods (squid, octopus, and cuttlefish) have evolved remarkable ability to manipulate light and alter their appearance via the combination of pigmentary and structural elements. Uppermost in the dermis, chromatophore organs proportionally regulate coloration by the stretching of pigmentary sacs [1; 2]. While the structural coloration is produced by two classes of cells called iridocytes and leucophores. Distributed on periphery of iridocytes, Bragg reflectors are periodically stacked lamellae which present iridescence by multilayer interference [3; 4]. Leucophores that contain granular vesicles are responsible for the production of bright white, by unselectively reflecting all incident light [5].

Interestingly, both periodically stacked lamellae in iridocytes [3; 4] and intracellular vesicles in leucophores [6; 7; 8] are unique and composed of reflectin proteins. Reflectin proteins are exclusively expressed in cephalopods, which members possess different number of canonical reflectin motifs in different sequential locations [3; 9; 10]. Guan et.al. reported that reflectin motifs may be traced to a 24-bp transposon-like DNA fragment from the symbiotic bioluminescent bacterium *Vibrio fischeri* [11]. Afterward, million years of self-replication and translocation of that transposon leads to the formation of prosperous reflectin family. For example, in *Doryteuthis. Opalescens*, as one representative of the most recently evolved Loliginid species, four distinct reflectins, reflectins A1, A2, B and C, have been identified and well characterized [3; 9; 10].

Numerous *in vitro* assays suggested that the dynamic condensation, folding and hierarchical assembly of reflectin proteins play fundamental roles during the formation and regulation of those structural coloration elements in squids [9; 10; 12]. Recently, Daniel E. Morse and his team have preliminarily defined reflectin proteins as a new group of intrinsically disordered proteins (IDPs), and suggested that transient liquid-liquid phase separation (LLPS) may dominate reflectin assembly [10]. The dynamic reflectin assembly properties and the intricate reflectin-based biophotonic systems have already inspired the development of various next-generation tunable photonic [13; 14; 15] and electronic platforms and devices [16; 17; 18].

In spite of impressive breakthroughs achieved by abovementioned works, knowledge about how reflectins molecules are spatially organized to form proteinaceous layers or droplets is still limited. Moreover, the limited existing information is mostly obtained from *in-tube* assays. It is necessary and interesting to explore reflecitns properties in cells.

To investigate their intracellular behaviors, we engineered HEK-293T cells to express either intact RfA1, RfA2, RfB1 or RfC. These proteins are the most widely studied reflectins, with different self-assembly characteristics [10; 12] and membrane-bound properties [19].

After transfection, reflectins were found to phase out from the crowded intracellular milieu, similar to what Atrouli [20] and Junko [21] described. Beyond that, more dramatic is the exclusive distribution of RfA1 particles in the cytoplasm, while spherical RfC droplets only existed in the nucleoplasm. As an intermediate state, RfA2 and RfB1 condensates distribute in both cytoplasm and nucleoplasm. This is notable evidence for their functional discrepancy, as well as their evolutionary process. Considering the difficulty of witnessing the molecular evolution that takes million years, to genetically design variants by cloning technology is much easier and operable. Therefore, we took the advantage of reflectins programmable sequences and designed a series of RfA1 truncations. By cutting off reflectin motifs (RMs) one by one from RfA1, the length and constitution of RfA1 variants are more and more similar to shorter natural reflectins (RfA2, RfB1 and RfC). Surprising but reasonable, as the RfA1 variants got shorter and shorter, protein droplets gradually change their localization from cytoplasm to nucleoplasm, making them comparable analogs to the shortest RfC.

This entrance of reflectins and variants into nucleus is a straightforward clue to delineate their intracellular transportation mechanism. Viewing from the vertical section, the pore at the bottom of Bragg lamellae approximately equals to the size of ciliary pore and nuclear pore. Updating evidence suggest that protrusive ciliary pore and nuclear pore share components and machinery [22; 23], including Ran GTPase, importins [24] and nucleoporins (Nups) [25]. Since RAN and Nups were identified to facilitate the entrance of RfA1 variants into nucleus in this study, it is reasonable to speculate that the entry of reflectins into lamellae follows the similar mechanism.

Moreover, an extensive interplay between RfA1 and cytoskeleton was also observed. Firstly, CoIP-MS survey indicated that RfA1 strongly bind to actin and actin binding proteins. Considering the essential role of actin filaments in formations of protrusive structures such as lamellipodia and ruffles [26; 27; 28; 29], this tight interaction between RfA1 and actin system suggests that RfA1 is capable to anchor at Bragg lamella, or even participate in the organization of laminar morphology. Secondly, RfA1 was detected to move towards the microtubules minus-ends and enrich in microtubule organizing center (MTOC), causing significant effects on spindle organization and cell division at transcriptomic level. As unpolarized cells, the microtubules minus-ends of HEK-293T are located at or around the centrosome, orienting the movement of RfA1 to MTOC. However, in epithelial cells, most MTs are non-centrosomal and align along the apico-basal polarity axis of the cell. Therefore, the enrichment of RfA1 around MTOC in this study is not ambivalent to its anchorage in Bragg lamellae. On the contrary, the MT minus-end-directed movement of RfA1 allows its recruitment beneath cytomembrane in iridocytes, storing the materials reserves and contributing to the construction of the delicate biophotonic reflectors.

## Results

### Phase separation and selective intracellular localization of reflectin proteins

RfA1, A2, and B1 possess different numbers and localization of two types of canonical reflectin motifs (Fig. 1a), the regular reflectin motifs (RMs, [M/FD(X)5MD(X)5MDX3/4]) and a N-terminal reflectin motif (RM_N_, [MEPMSRM(T/S)MDF(H/Q)GR(Y/L)(I/M)DS(M/Q)(G/D)R(I/M)VDP(R/G)]) (Fig. 2a). Specifically, RfC is the shortest reflectin, contains a GMXX motif and RM*. The GMXX motif is a unique region of increased overall hydrophobicity composed of a four amino acid repeat, where ‘X’ represents less conserved locations within the repeat. Asterisk marked RM* of RfC contains substantial deviations in sequence not observed in any other reflectin motifs [30] (Fig. 2b). The disorder tendencies of reflectins were calculated by PONDR [31], DISOclust [32; 33], ESpritz[34]. The possibility to form protein droplets via liquid-liquid phase separation is also predicted by FuzDrop [35]. These calculations indicate high disorder (Supplementary Fig. 1) and LLPS tendency (Supplementary Fig. 2) of four reflectin proteins. To verify these computational calculations, reflectin proteins were introduced into HEK-293T cells.

**Fig.1.**
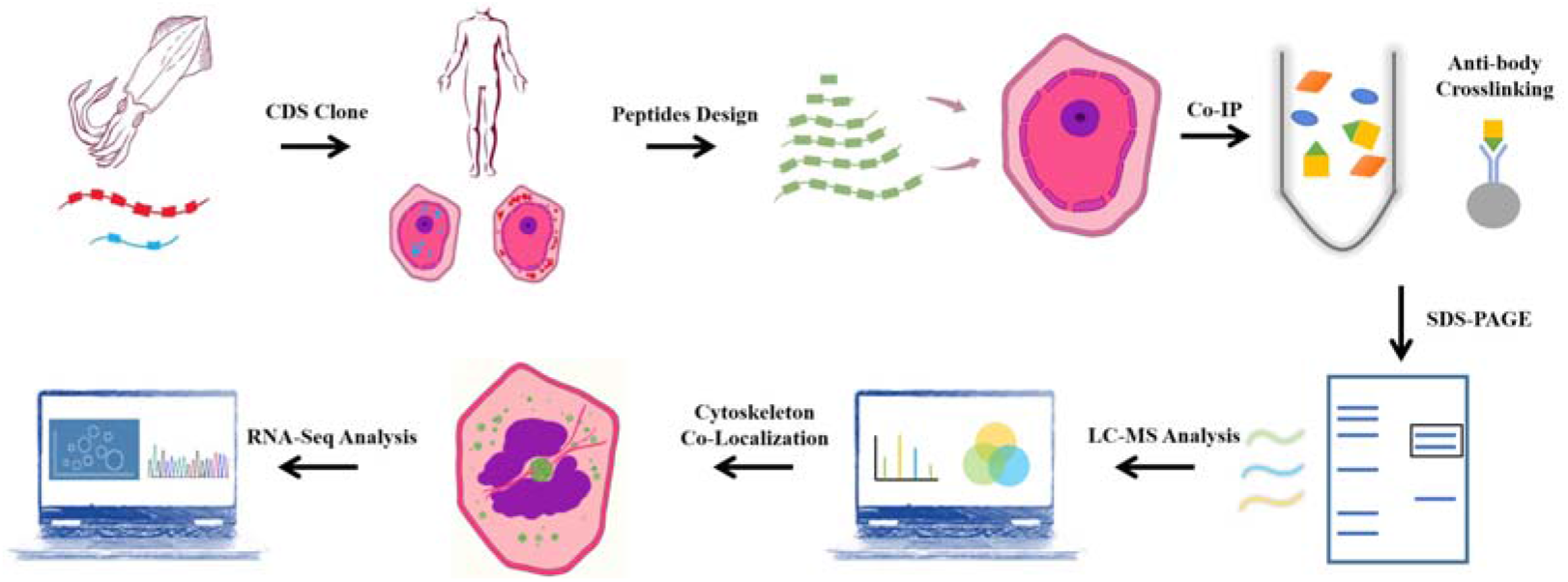
Overview of the workflow. CDS of squids reflectin proteins were obtained from NCBI, optimized according to mammal cell expression preference and introduce into HEK-293T cells by lipofection. RfA1 truncated variants were designed and cloned into pEGFP-C1 vectors. Transfected cells were imaged by confocal microscope. Anti-GFP monoclonal antibody was used to pull down GFP-tagged RfA1 variants and their binding proteins. The immunoprecipitates were separated using SDS-PAGE, and gel lanes were digested with trypsin for LC-MS/MS analysis. Co-localization of RfA1 and actin cytoskeleton was confirmed by confocal microscope. RNA-Seq was employed to globally investigate the influence of RfA1 expression on cells.

**Fig.2.**
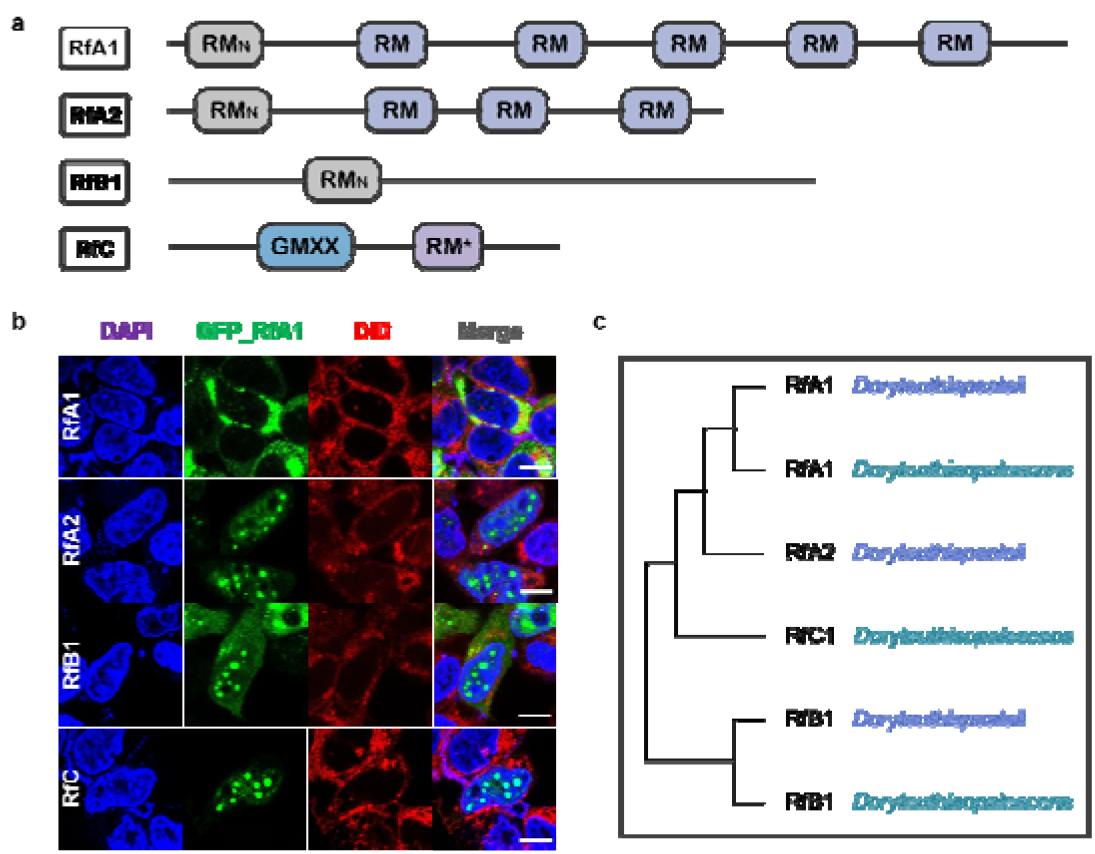
Transfection of reflectins induces protein condensates formation in selective intracellular regions. **a** Schematic Reflectins sequences. Reflectin Motifs are designated by boxes, while reflectin linkers (RLs) are lines. **b** Confocal images of GFP_Reflectin condensates in HEK-293T cells. Nuclei and membrane were stained with DAPI (blue) and DiD (red) respectively, Scale bars 10μm. **c** cladogram of RfA1, RfA2, RfB1 and RfC.

After transfection and culturing for 12hr, all RfA1, RfA2, RfB1, RfC phased out from the cytoplasm or nucleoplasm, which is consistent with computational predictions. However, their difference is more significant. For RfA1, protein condensates exclusively enriched in the cytoplasm. As a sharp contrast, the perfect spherical RfC droplets only locate in the nucleoplasm. While the distribution of RfA2 and RfB1 seems to stay in a middle state: spherical protein droplets mainly existed in the nucleus, with the minority stayed in the cytoplasm.

According to Guan et al. report, reflectin genes in cephalopods come from a transposon in symbiotic *Vibrio fischeri* [11]. The repetitions of conserved motifs in reflectin sequences could be the results of transposon self-replication and translocation. Thus, it is reasonable to speculate that reflectins with different motifs stay at different evolution stages, while shorter ones with simpler sequence structure should be ancestors.

View from this point, RfC with only a GMXX motif and RM* should be the mother protein, while the RfA1 with a RM_N_ and five RMs is the daughter protein. Based on this consideration, we started to construct truncated variants by cutting off the RMs from the RfA1 one by one, to make them compositional comparable to shorter natural reflectins. The dynamic intracellular performance of these RfA1 variants was then investigated.

### Intracellular localization preferences of RfA1 variations

Six pairs of primers were designed to clone RfA1 truncations (Supplement Table. 1). The PCR products responsible for coding of six peptides, including RM_N_, RM_Nto1_, RM_Nto2_, RM_Nto3_, RM_Nto4_, and RM_Nto5_ (schematics shown in Fig. 3a), were subsequently ligated to the vector pEGFP-C1. After transfection, the expressions of GFP-tagged RfA1 truncations were validated by anti-GFP Western blotting (Fig. 3b). The molecular weights of truncations were calculated by ExPASy ProtParam tool and confirmed by Western blotting.

**Fig.3.**
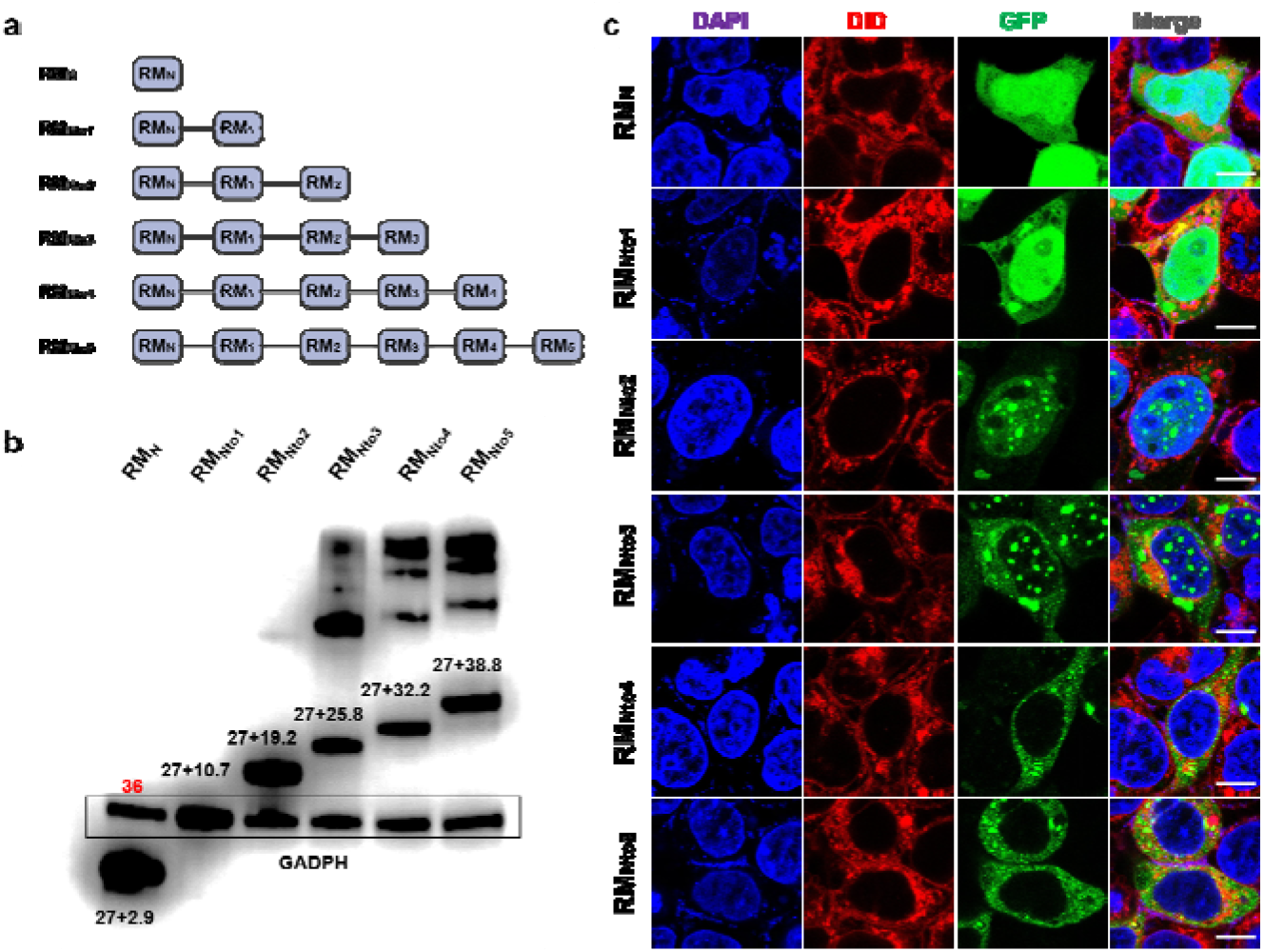
Expression of RfA1 variants in HEK293T cells. **a** schematics of RfA1 truncated variants. **b** the expression of RfA1 variants were validated by Western blotting, indicated by anti-GFP antibodies. Molecular weight of EGFP is 27 kDa; GADPH is 36 kDa. **c** phase separation and cyto-/nucleo-localization preferences of truncated RfA1 derivatives. Nuclei and membrane are stained with DAPI (blue) and DiD (red), RfA1-derived peptides are indicated by tandem EGFP (green), Scale bars 10μm.

To investigate the intracellular localization, GFP-tagged RfA1 truncations was observed by confocal image system (Fig. 3c). Nuclei and membrane are stained with DAPI (blue) and DiD (red), RfA1-derived peptides are indicated by tandem EGFP (green). For cells transfected with pEGFP-C1-RM_N_ and pEGFP-C1-RM_Nto1_, though the peptide condensation seems absent, a remarkable enrichment of green fluorescence signal in nucleus areas appeared. Amazingly, this nucleus-enrichment is executed more thoroughly by RM_Nto2_, along with the recurrence of proteinaceous droplets. Ulteriorly increase of conserved motifs, RM_Nto3_ are found to distribute in both cyto- and nucleo-plasm. For longer derivatives with more motifs, condensates of RM_Nto4_ and RM_Nto5_ were blocked out from nuclei and located exclusively in cytoplasmic localization, extremely resemble their template RfA1.

Briefly, RM_Nto2_ is most similar to RfC among all variants, which forms condensates exclusively in nucleoplasm. More significantly, RM_Nto2_ is also a definte boundary: truncations shorter than RM_Nto2_ tend to enrich in nuclei, but cannot form droplets; derivatives longer than RM_Nto2_ start to escape from the nuclei, accumulate and condensate into droplets in cytoplasm.

Also noteworthy are smears in PAGE lanes of RM_Nto3_, RM_Nto4_, and RM_Nto5_ in Fig. 3b. These are not sample contaminations or non-specific labeling. Subsequent CoIP-MS results reveal various cellular components tightly interact with RfA1 variants, which assistant their phase separation and diverse cellular localization.

### Identification of the RfA1/RM_Nto2_ binding proteins

To explore the mechanism underlying phase separation and distinguished distribution of RfA1 and RM_Nto2_, a monoclonal antibody of GFP was prepared and a CoIP-MS approach was used to identify protein-protein interactions (PPIs).

Aforementioned, there must be a massive interaction between RfA1 variants and host cell proteins, resulting in the smears presented in Fig. 3b. Therefore, valuable information will be lost if only one single band was collected and analyzed. However, mixed samples consist of multiple bands inevitably bring in meaningless miscellaneous proteins. Based on these considerations, cells expressed RM_N_ was selected as a control (Fig. 4a), since RM_N_ presented no significant phase separation or selective distribution behavior.

**Fig.4.**
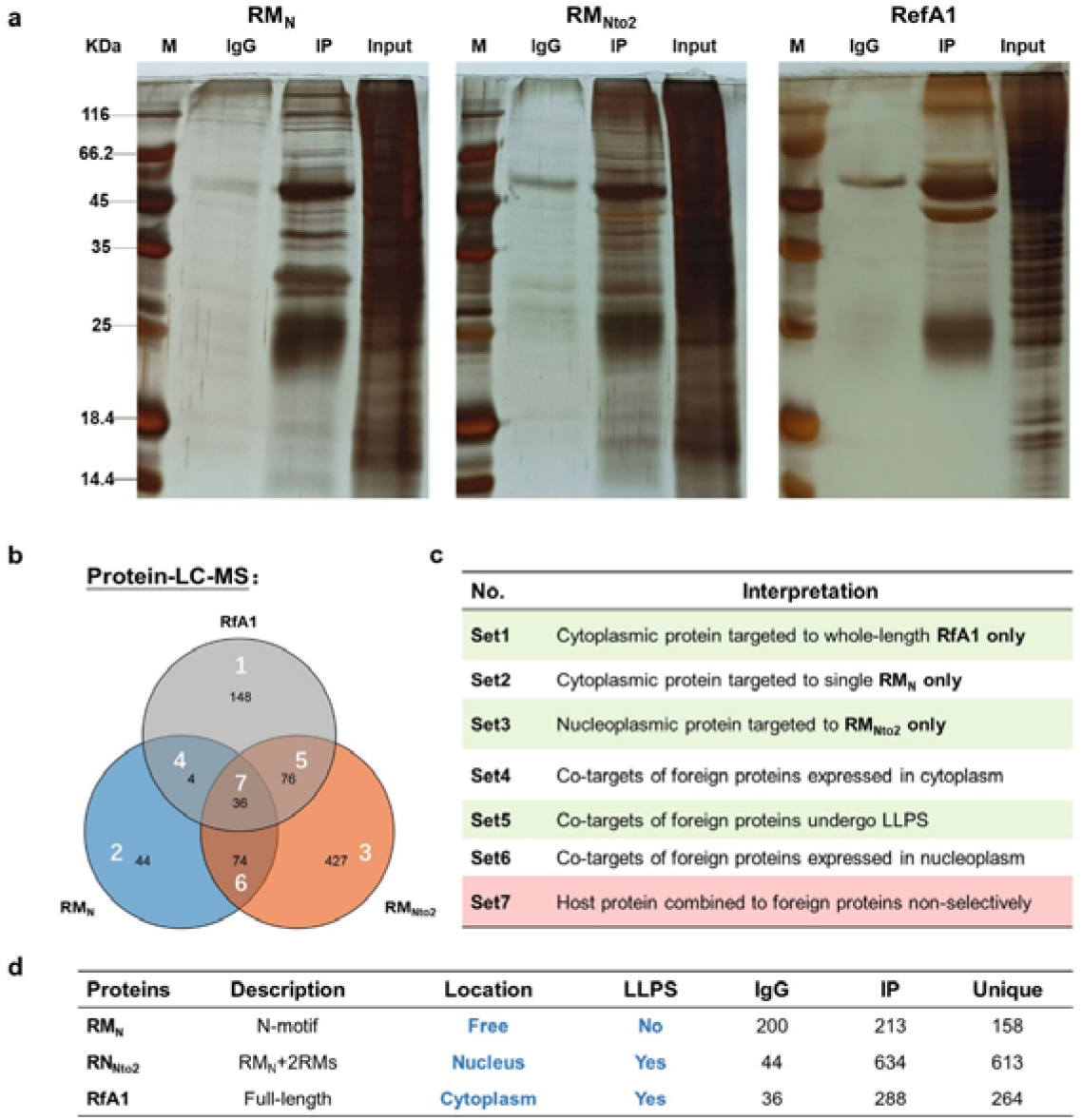
Identification of interaction proteins targeted to RfA1 and RM_Nto2_. **a** silver staining of IP products, whole IP bands were sent to LC-MS. **b** the intersection of three LC-MS results was shown in the Boolean graph. **c** interpretations of areas in the Boolean graph. **d** PPI summary. Unique(RM_Nto2_)=IP(RM_Nto2_) - IP(RM_Nto2_)⋂IgG (RM_Nto2_) - IP(RM_N_)⋂IgG (RM_N_); Unique(RfA1)= IP(RfA1) - IP(RfA1)⋂IgG (RfA1) - IP(RM_N_)⋂IgG (RM_N_).

As shown in the Boolean graph (Fig. 4b), overlays Set4/6/7 will be eliminated as backgrounds. Within Set1 are the proteins combined to whole-length RfA1, and contribute to the phase separation and cytoplasmic localization; proteins in Set3 promote the importation of RM_Nto2_ into nuclei and its phase out from the nucleoplasm; co-targets of RM_Nto2_ and RfA1 are sorted into Set 5 (Fig. 4c). Protein-protein interactions are summarized in Fig. 4d.

Gene Ontology (GO) annotation analysis was then performed to investigate biological processes associated with RfA1 and RM_Nto2_ interacting proteins. Items of interests with adjusted p-value < 0.05 are profiled and shown in Fig. 5 (Gene lists in Supplement Table. 2, PPI Networks in Supplement Fig. 3).

**Fig.5.**
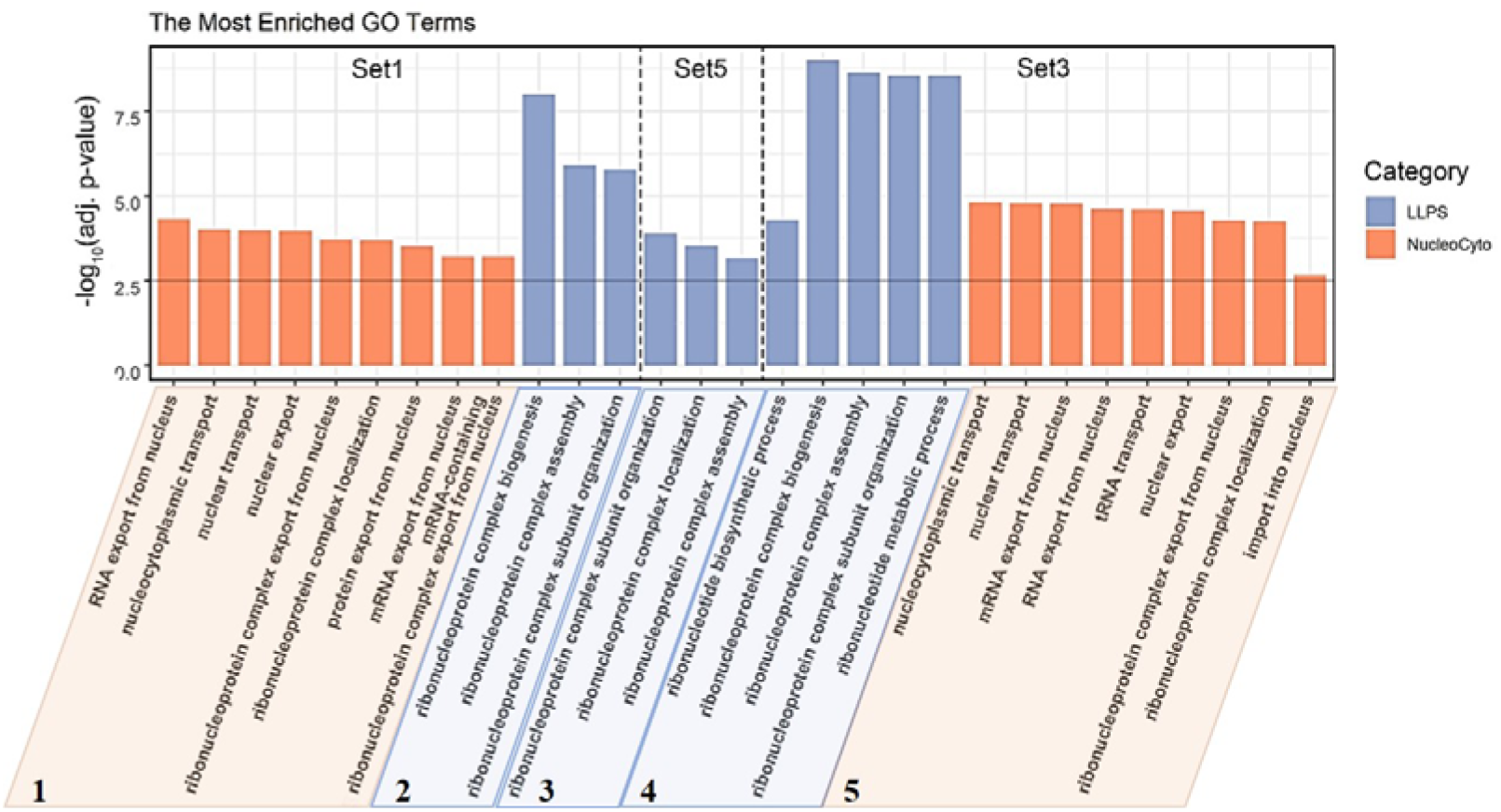
GO biological processes annotation of proteins associated with LLPS and cellular localization, in Set1, Set5 and Set3. Gene list in Set1, Set5 and Set3 were converted to ENTREZ ID and R package clusterProfiler were used to perform Gene ontology annotation analysis. LLPS and cellular localization-associated BP terms were showed in the figure, the threshold was set as log_10_(adjusted p-value) < −2.5.

Take Set1-1 for example, since RfA1 bind to proteins like HNRNPA1 [36; 37], SRSF1 [38] and RAN [39; 40] and participate in process including “protein export from nucleus”, it is comprehensible that the RfA1 condensates are exclusively distributed in the cytoplasm. Similarly, RANBP2 [41] and various NUPs [42] in Set3-5 facilitate the importation of RN_Nto2_ into nuclei. Besides, it is also valuable evidence to interpretate the transportation of reflectins into iridocytes Bragg lamellae. Viewing from the vertical section, the pore at the bottom of Bragg lamellae has a similar size to a ciliary pore. Recent studies indicate that ciliary pore and the nuclear pore share components and machinery [22; 23]. Ran GTPase, importins [24] and nucleoporins (Nups) [25] were found to involve in protein transportation across both ciliary pore and nuclear pore. Here, RAN and Nups were identified to mediate the access of RfA1 variants into nucleus. It is presumable that the entry of reflectins into Bragg lamellae follows the alike mechanism, under the guidance of similar components (RAN, Nups, etc.).

One remarkable common points of Set1-2, Set5-3 and Set3-4 is the involvement of “ribonucleoprotein complex” or “ribonucleotide metabolic”. The interaction with ribonucleoprotein complexes brings in a massive interplay with proteins and mRNA [43; 44], leads to an almost inevitable phase separation. This is consisted with results in Fig. 3c. However, the involvement of ribonucleoprotein complex makes the situation more elusive. It is equivocal whether RfA1/RN_Nto2_ molecules gathered together and phase out from the surrounding by their own, or they were drawn into the formation processes of ribonucleoprotein complex bodies.

Moreover, PPI information also demonstrates that RfA1 extensively interacts with actin and actin binding proteins (ABPs) (Fig. 6, Set1-1, Gene lists in Supplement Table. 3, PPI Networks in Supplement Fig. 4). These proteins are in charge of various biological processes, including actin polymerization, nucleate assembly of new filaments, promote elongation [45; 46]. Although there is a huge difference between HEK-293T cells and squids iridocytes, the evolutionarily conservative of actins among algae, amoeba, fungi, and animals [47; 48] eliminated this drawback. Besides, considering the critical role of actin filaments in formations of protrusive structures such as lamellipodia and ruffles [26; 27; 28; 29], results presented here highly suggest RfA1’s potential contribution in the establishment and regulation of Bragg lamellae via its collaboration with actin system. By cooperating with actin and ABPs, RfA1 is able to anchor at the inside of Brag lamella, or even participate in the organization of the actin network and hereby regulates the cell morphogenesis.

**Fig.6.**
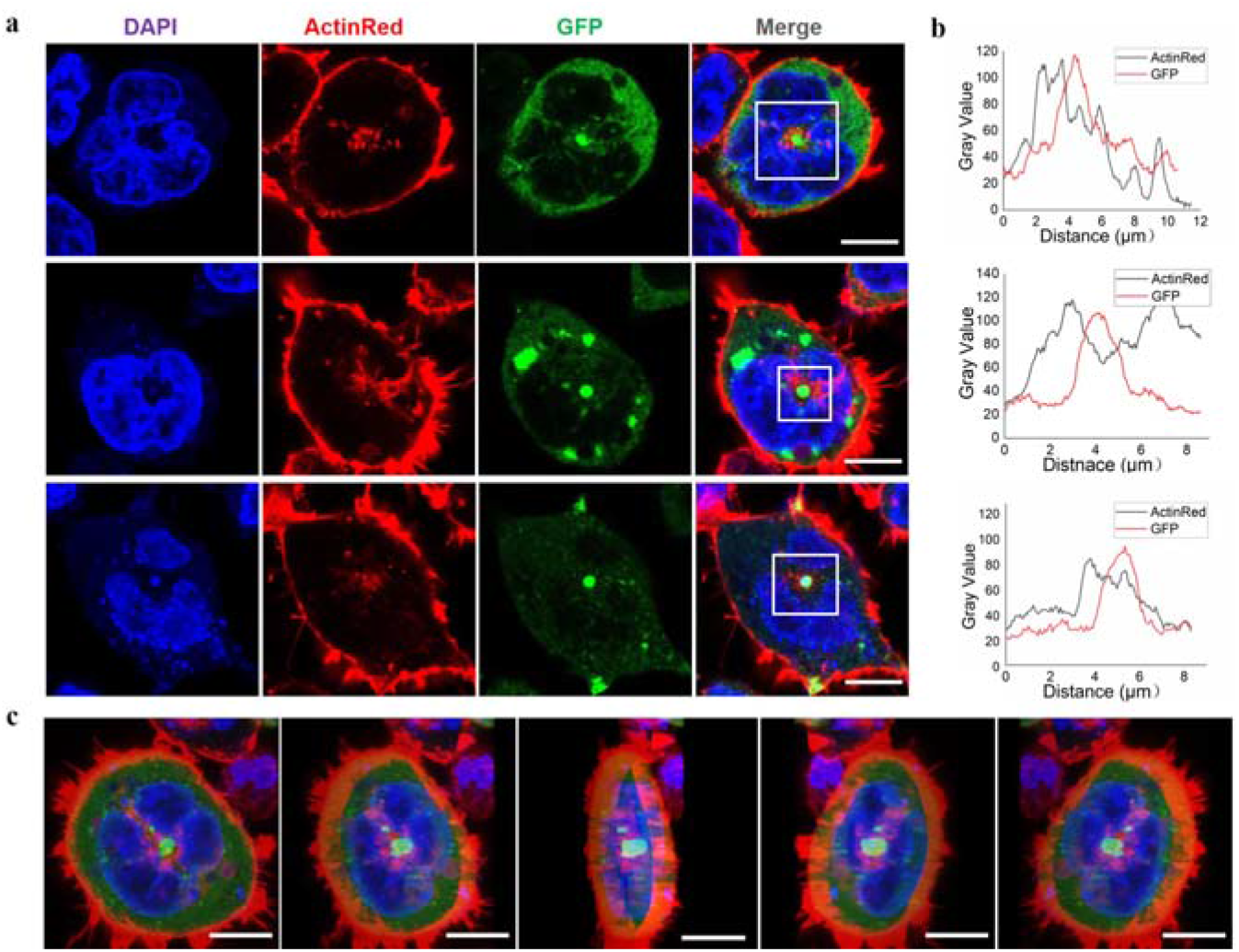
Appearance of spherical RfA1 condensates in the center or on the central axis of transfected cells. **a** RfA1 condensates located in the central or on the central axis of cells. Nuclei and actin are stained with DAPI (blue) and ActinRed 555 (red), RfA1 are indicated by tandem EGFP (green), Scale Bars 10μm. **b** statistics of fluorescence signal distribution in selected areas. **c** 3D images of a transfected cell, with a green RfA1 condensates located in the center, Scale Bars 10μm (Supplement Movie. 2).

### Verification of co-localization of RfA1 and actin system

Actin should be indispensable for either the establishment or the regulation of delicate Bragg reflectors in iridocytes. However, so far, except for the Co-IP results presented in this work, no evidence has been issued about the interaction between reflectin proteins and the cytoskeleton. As a preliminary start, we stained the HEK-293T cells with ActinRed 555 (Thermo) to trace the actin network, and to verify the potential relationship between RfA1 and cytoskeleton.

One notable trouble is the ubiquity of the actin cytoskeleton. It is hard to judge the co-localization of RfA1 and actin (Supplement Movie. 1). However, after viewing thousands of cells, an interesting scenario began to emerge. For plenty of transfected cells, there is a bright green sphere in the center or on the central axis, which are condensates of RfA1 (Fig. 6a, GFP channel). This central localized protein condensates were not found in RfA2, RfB1 or RfC transfected cells. Around these spherical condensates, the red signal of stained actin is also significantly stronger than other areas (Fig. 6a, ActinRed channel). Fluorescence statistics of selected areas in Fig. 6b shown an almost synchronized fluctuation of ActinRed and GFP signals, indicating their co-localization (Raw data in Supplement Table.4).

### RfA1 interacts with microtubules and moves minus-end orientated

Except for the fluorescent staining assays, RNA-seq results also strongly suggested the interaction between RfA1 and the cytoskeleton. Cells transfected with pEGFP-C1-RfA1 and no-load pEGFP-C1 (set as control) were harvested for sequencing. After Metascape screening [49], 461 differentially expressed unigenes were identified in RfA1 expressed cells (Supplement Table.5). Gene Ontology (GO) analysis was then performed to annotate differentially expressed genes (DEGs). Selected items of interest are listed in Fig. 7a. Genes evolved in biological processes such as “regulation of microtubule cytoskeleton organization”, “microtubule polymerization or depolymerization”, “microtubule nucleation” were found to respond to RfA1 expression. Correspondingly, these microtubule relevant genes were also sorted to cellular component items, including “spindle microtubule”, “spindle pole centrosome”, “mitotic spindle”, “spindle” [50; 51], which indicates the tight relationship between RfA1 and microtubules.

**Fig.7.**
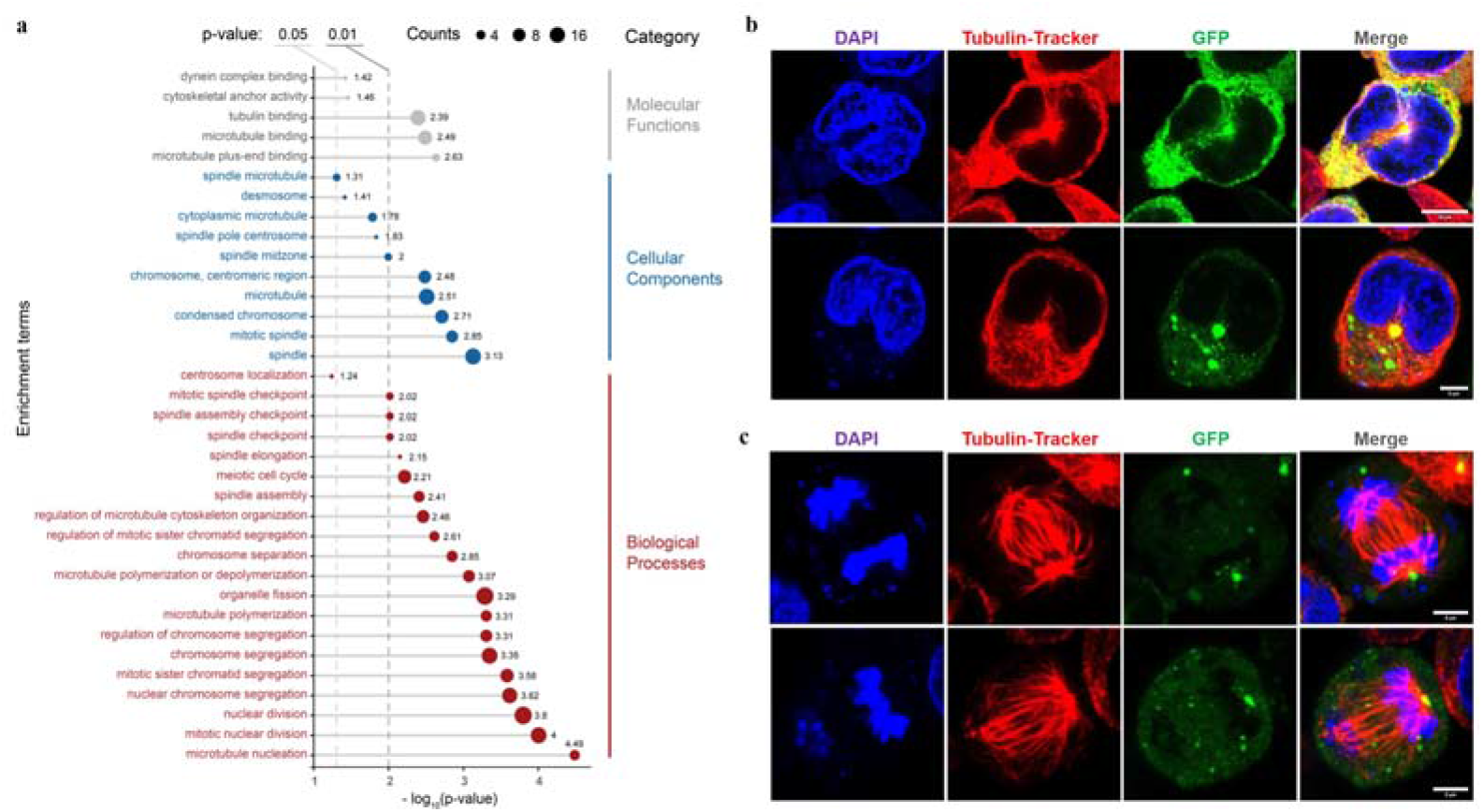
Effects of RfA1 on microtubule and mitosis. **a.** GO annotation of DEGs between pEGFP-C1 transfected cells and pEGFP-C1-RfA1 transfected cells. Genes were converted to ENTREZ ID and R package clusterProfiler were used to perform Gene ontology annotation analysis. Biological Process, Cellular Component and Molecular Function terms were showed in the figure, the threshold was set as p-value < 0.05. **b.** RfA1 co-localization with microtubule networks in the center, in interphase cells. Nuclei and microtubule are stained with DAPI (blue) and Tubulin-Tracker Red (Beyotime), RfA1 are indicated by tandem EGFP (green), Scale Bars 10μm. **c.** RfA1 located at the spindle center, in late mitosis cells. Nuclei and microtubule are stained with DAPI (blue) and Tubulin-Tracker Red (Beyotime), RfA1 are indicated by tandem EGFP (green), Scale Bars 5μm (Supplement Movie. 3,4).

The analysis based on RNA-Seq was further confirmed by confocal observation. After staining the fixed cells with Tubulin-Tracker Red (Beyotime), RfA1 condensates were significantly enriched at MTOC, located around centrosome (Fig. 7b). For cells stays in late mitosis, RfA1 condensates were transported along the microtubules, chasing the movements of daughter centrosomes (Fig. 7c, Supplement Movie. 3,4). The obvious phenomena here is the spatial enrichment of RfA1 at MOTC, which can be severely abolished by nocodazole treatment (data not shown). What less eye-catching but equally important is the microtubule minus-end-directed trafficking of RfA1. For HEK-293T cells, microtubule minus-ends are located at or around the centrosome, guiding the movement of RfA1 to MTOC (Fig. 7b). However, in polarized epithelial cells, microtubules are oriented apicobasally, with their minus-ends directed toward the apical side of the cells [52]. In these cases, the microtubules minus-ends can be re-orientated and anchored to the cytomembrane [53]. That is to say, in epithelial cells including iridocytes, microtubules system is able to transport reflectins towards Bragg lamellae which protrude out from cells like specialized cilia, fulfilling the material requirements for the establishment of biophotonic reflectors.

## Discussion

In summary, by taking advantage of reflectins programmable sequences, we realized the expression of four natural reflectins (RfA1, RfA2, RfB1 and RfC) and six engineered RfA1 in cells, and found their tunable intracellular localization and extensive interaction with cytoskeleton. We believe there are multiple reasons to support the significance of this work.

Firstly, the cyto-/nucleoplasmic localization preference presented here could be evidence for the molecular evolution of reflectin proteins. According to Guan’s report, reflectin genes in cephalopods come from a transposon in symbiotic *Vibrio fischeri* [11]. Afterward, million years of replication and translocation of this transposon results in the formation of reflectins with different length, different motifs and different properties [3; 9; 10; 54]. In this study, RfA1 consists of one RM_N_ and five RMs exclusively enriched in cytoplasm, while RfC contains only one GMXX and a RM* stayed in nucleus. By truncating RfA1 gradually, shorter peptide derivatives (e.g., RM_Nto2_) started to localize to nucleus, behaved exactly similar to RfC. In this point of view, shorter and simpler RfC should originally emerge as the ancestor molecular in nucleus. Then longer reflectins began to be assembled and escape from the nucleus into cytoplasm (Fig. 8a).

**Fig.8.**
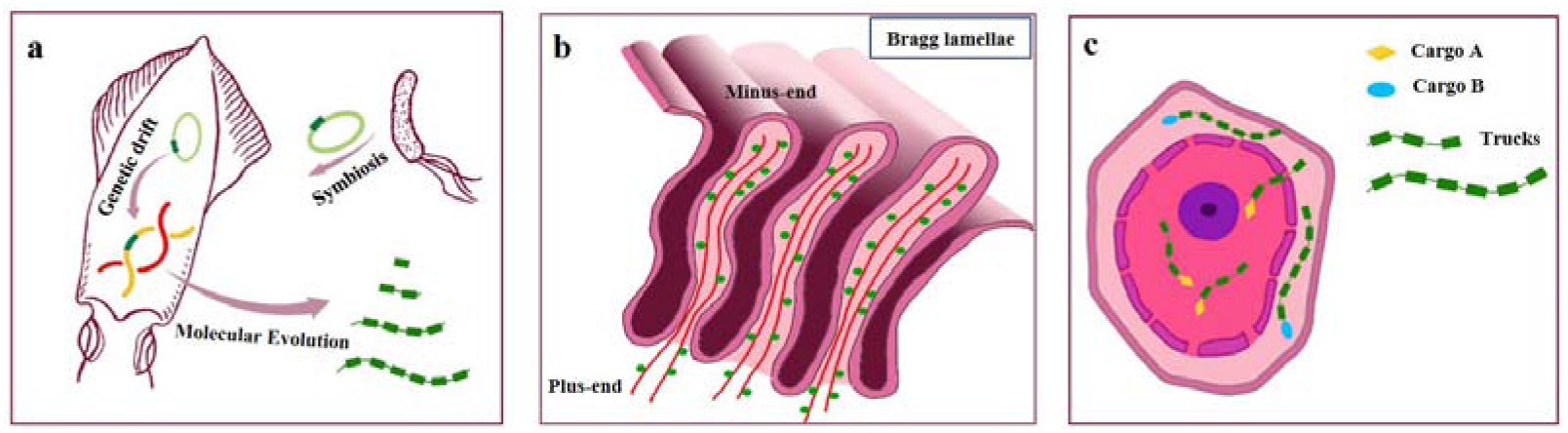
Models. **a** origin and evolution of reflectin proteins. **b** Microtubule minus-end-directed transportation of reflectin. **c** schematic for RfA1 derivatives as programmable and oriented cargo carriers.

Secondly, the nucleus entry of short reflectins and microtubule minus-end-orientated RfA1 transportation delineate their potential organization mode in iridocytes, which lay the material foundation for Bragg lamellae establishment. 1) Seeing from longitudinal section, the pore at the bottom should be in charge of material transportation into and out of lamellae(Fig. 8b). Though knowledge about protein transportation between cytoplasm and Bragg lamellae is absent, the entrance of RfA1 truncations and short reflectins into nucleoplasm is clearly observed. Considering the emerging hypothesis that the pore structures (such as ciliary pore, nuclear pore) share components and machinery [22; 23; 24; 25], the transportation of RfA1 into lamellae may follow the similar approaches. 2) Both CoIP-MS analysis and RNA-seq survey revealed the extensive interactions between RfA1 and cytoskeleton. Particularly, RfA1 was observed to enrich at MTOC in late mitosis cells, as a result of microtubules minus-end-directed transportation. In epithelial cells such as iridocytes, since the microtubules are re-oriented apicobasally [52; 53], the direction of reflectins movements can be inverted and points to cytomembrane, allowing the protein accumulation for Bragg lamellae construction (Fig. 8b).

Thirdly, reflectins are natural block copolymers, as well as programmable biomaterial. Their intracellular localization could be easily adjusted by the tunable sequence length. If we take GFP as a molecular cargo, reflectin derivatives can be regarded as intelligent vehicles to transport cargoes to pre-selected destinations (e.g., cytoplasm or nucleus, Fig. 8c). On the other hand, reflectin proteins or reflectin variants described in this study make up a powerful arsenal to engineer human cells to achieve coloration, as Atrouli [20] and Junko [21] brilliant works have demonstrated.

## Methods

### Construction of recombinant pEGFP-C1 vectors

Nucleotide sequence of *D. (Loligo) pealeii* reflectin A1 (RfA1) (Genbank: ACZ57764.1) and *D. (Loligo) Opalescens* reflectin C (Genbank: AIN36559.1) were optimized for human cell expression, then synthesized and sequencing-identified by Sangon Biotech®(Shanghai, China) Primers (F-GAATTCTAT GAATAGATATTTGAATAGACA; R-GGATCCATACATATGATAATCATAATA ATTT) were designed to introduce *EcoR I* and *BamH I* cutting sites, so the modified RfA1 CDS can be constructed into pEGFP-C1 via a standard restriction enzyme cloning process. As for truncated RfA1 derivatives, six pairs of primers were coupled used. Take this for example, if RM_N_-F and RM_3_-R primers were selected, then a nucleotide sequence responsible for the coding of RM_N_-RL_1_-RM_1_-RL_2_-RM_2_-RL_3_-RM_3_ (which is simplified as RM_Nto3_ in this paper) will be obtained after PCR. In the meanwhile, 5′ GCATGGACGAGCTGTACAAG 3′ and 5′ TTATGATCAG-TTATCTAGAT 3′ were added to those F-primers and R-primers respectively during primers synthesis, which enables sequences to be ligated to pEGFP-C1 by Ready-to-Use Seamless Cloning Kit from Sangon Biotech®(Shanghai, China).

### Growth and transfection of human cells

HEK-293T cells (ATCC®, CRL-3216^TM^) were cultured on plastic dishes in Dulbecco’s Modified Eagle Medium (DMEM, Gibco^TM^) supplemented with 10% fetal bovine serum (FBS, Gibco^TM^) at a temperature of 37 °C and under 5% CO_2_. One day before transfection, cells were seeded at ~33% of the confluent density for the glass bottom dishes from Cellvis (California, USA), and grown for another 24 h. Then transfection mixture containing Lipofectamine 3000 (Thermo Scientific) and recombinant vectors was added to the medium, incubating for ~16 to ~24 h. For CCK-8 tests, 1×10^4^ cells were seeded into each hole of 96-well plates one day before transfection, then transfected with recombinant vectors and incubated for another 24 h. After that, 10μl of CCK-8 solution will be added into wells for a ~2 to ~4 h chromogenic reaction. OD_450_ was detected by Multiskan FC (Thermo Scientific).

### Fluorescence microscopy of stained cells

Transfected HEK-293T cells grown in Cellvis plastic dishes were firstly fixed with 4% paraformaldehyde at room temperature for 30 mins, then stained with DiD or ActinRed (diluted in 0.5% triton X-100 PBS) for ~30 mins after PBS rinses. After washing off the fluorescent dye with PBS, fixed cells will be embedded in DAPI-Fluoromount (Beyotime, Shanghai, China) and characterized with a Leica TCS SP8 imaging system in fluorescence imaging mode. The resulting images were analyzed with ImageJ (Java 1.8.0_172/1.52b) [55].

### Online computational prediction tools

Disorder prediction of RfA1 and derivatives were conducted with PONDER®(http://www.pondr.com/), DISOclust (http://www.reading.ac.uk/bioinf-/DISOclust/) and ESpritz (http://protein.bio.unipd.it/espritz/). The probability undergoes liquid-liquid phase separation was calculated by FuzPred (http://protdyn-fuzpred.org/)

### SDS-PAGE, Western Blotting

Total proteins of cells were extracted by Immunoprecipitation Kit (Sangon). Protein samples were separated in SDS-PAGE and transferred to polyvinylidene fluoride (PVDF) filter membranes (Millipore, USA) for immune-hybridization. After 1 hour blocking in PBST (phosphate buffered saline containing 0.05% Tween-20 and 2.5% BSA), the membranes were incubated with one of the following primary antibodies with corresponding concentration: anti-GFP mouse monoclonal antibody (Sangon), anti-GAPDH mouse monoclonal antibody (Sangon). Subsequently, band visualization was performed by electro-chemiluminescence (ECL) and detected by Digit imaging system (Thermo, Japan).

### In-gel digestion

For enzymolysis in protein gel (gel strip, protein solution, protein freeze-dried powder), using a clean blade to dig out the strip of interest and cut it into 0.5-0.7 mm cubes. After decolorizing with test stain/silver staining decolorizing solution, wash the rubber block with 500μL of acetonitrile solution three times until the rubber particles are completely white. Add 500 μL of 10 mM DTT, water bath at 56°C for 30 min, low-speed centrifugation for reduction reaction; add 500 μL of decolorizing solution, mix at room temperature for 5-10 min, wash the gel spots once, centrifuge at low speed to discard the supernatant; quickly add 500 μL 55 mM IAM, placed in a dark room at room temperature for 30 minutes; centrifuge at low speed for alkylation reaction, then add 500 μL of decolorizing solution, mix at room temperature for 5-10 minutes, wash the gel spots once, centrifuge at low speed, and discard the supernatant; then add 500 μL of acetonitrile until the colloidal particles are completely white, vacuum dry for 5 min. Add 0.01 μg/μl, trypsin according to the volume of the gel, ice bath for 30 min, add an appropriate amount of 25 mM NH_4_HCO_3_ in pH8.0 enzymatic hydrolysis buffer, and enzymatic hydrolyze overnight at 37°C. After the enzymolysis is completed, add 300 μL extract, sonicate for 10 min, centrifuge at low speed, collect the supernatant, repeat twice, combine the obtained extracts and vacuum dry.

### Zip-tip Desalting

Dissolve the sample with 10-20 μL 0.2% TFA, centrifuge at 10,000 rpm for 20 min, and wet the Ziptip 15 times with a wetting solution. Equilibrate with a balance solution 10 times. Inhale the sample solution for 10 cycles. Blow 8 times with rinse liquid. Note: To avoid cross-contamination, aliquot 100 μL of flushing solution per tube, and use a separate tube for each sample. Add 50 μL of eluent to a clean EP tube, and pipette repeatedly to elute the peptide. Drain the sample.

### LC-MS/MS Analysis

The peptide samples were diluted to 1 μg/μL on the machine buffet. Set the sample volume to 5μL and collect the scan mode for 60 minutes. Scan the peptides with a mass-to-charge ratio of 350-1200 in the sample. The mass spectrometry data was collected using the Triple TOF 5600 +LC/MS system (AB SCIEX, USA). The peptide samples were dissolved in 2% acetonitrile/0.1%formic acid, and analyzed using the Triple TOF 5600 plus mass spectrometer coupled with the Eksigent nano LC system (AB SCIEX, USA). The peptide solution was added to the C18 capture column (3 μm, 350 μm×0.5 mm, AB Sciex, USA), and the C18 analytical column (3 μm, 75μm×150) was applied with a 60 min time gradient and a flow rate of 300 nL/min. mm, Welch Materials, Inc) for gradient elution. The two mobile phases are buffer A (2% acetonitrile/0.1%formic acid/98% H_2_O) and buffer B (98% acetonitrile/0.1% formic acid/2% H2O). For IDA (Information Dependent Acquisition), the MS spectrum is scanned with anion accumulation time of 250 ms, and the MS spectrum of 30 precursor ions is acquired with anion accumulation time of 50 ms. Collect MS1 spectrum in the range of 350-1200 m/z, and collect MS2 spectrum in the range of 100-1500 m/z. Set the precursor ion dynamic exclusion time to 15s.

Genes identified from LC-MS and RNA-Seq were converted to ENTREZ ID and R package clusterProfiler were used to perform Gene ontology annotation analysis [56]. The mass spectrometry proteomics data have been deposited to the Proteome Xchange Consortium via the PRIDE [57] partner repository with the dataset identifier PXD031350. RNA-Seq data has been uploaded to NCBI GEO database, numbered as GSE186861.

All data needed to evaluate the conclusions are present in the paper and/or the supplementary information. All other relevant data are available from the authors upon reasonable request.

## Abbreviations

reflectin: (Rf)
N-terminal reflectin motif: (RM_N_)
reflectin motifs: (RMs)
reflectin linkers: (RLs)
intrinsically disordered proteins: (IDPs)
liquid-liquid phase separation: (LLPS)
protein-protein interactions: (PPIs)
Gene Ontology: (GO)
microtubule: (MT)
microtubule organizing center: (MTOC)

## Acknowledgements

We acknowledge Prof. X. Zhu (SIBCB, CAS) and Prof. L. Qiu (MBL, HNU) for valuable discussion. We are grateful to the National Natural Science Foundation of China (Grants 31971291) and Natural Science Foundation of Hunan Province (Grants S2020JJQNJJ1684) for their financial support. J. Song gives his personal thanks to Prof. D.E. Morse and R. Levenson (MCDB, UCSB), for their devoted guidance from 2016 to 2018.

## Author contributions

J.S. designed and performed experiments, analyzed data, and completed the manuscript. C.L designed and performed experiments, analyzed the LC-MS data. L.Z. assisted in directing graduate students to carry out experiments. B.L. and L.L. performed experiments. Z.Y. analyzed data. T.M. designed and plotted the schematic diagrams in Fig. 1 and Fig. 8. W.W. and B.H. designed experiments and wrote the manuscript.

---

**Fig.1.**
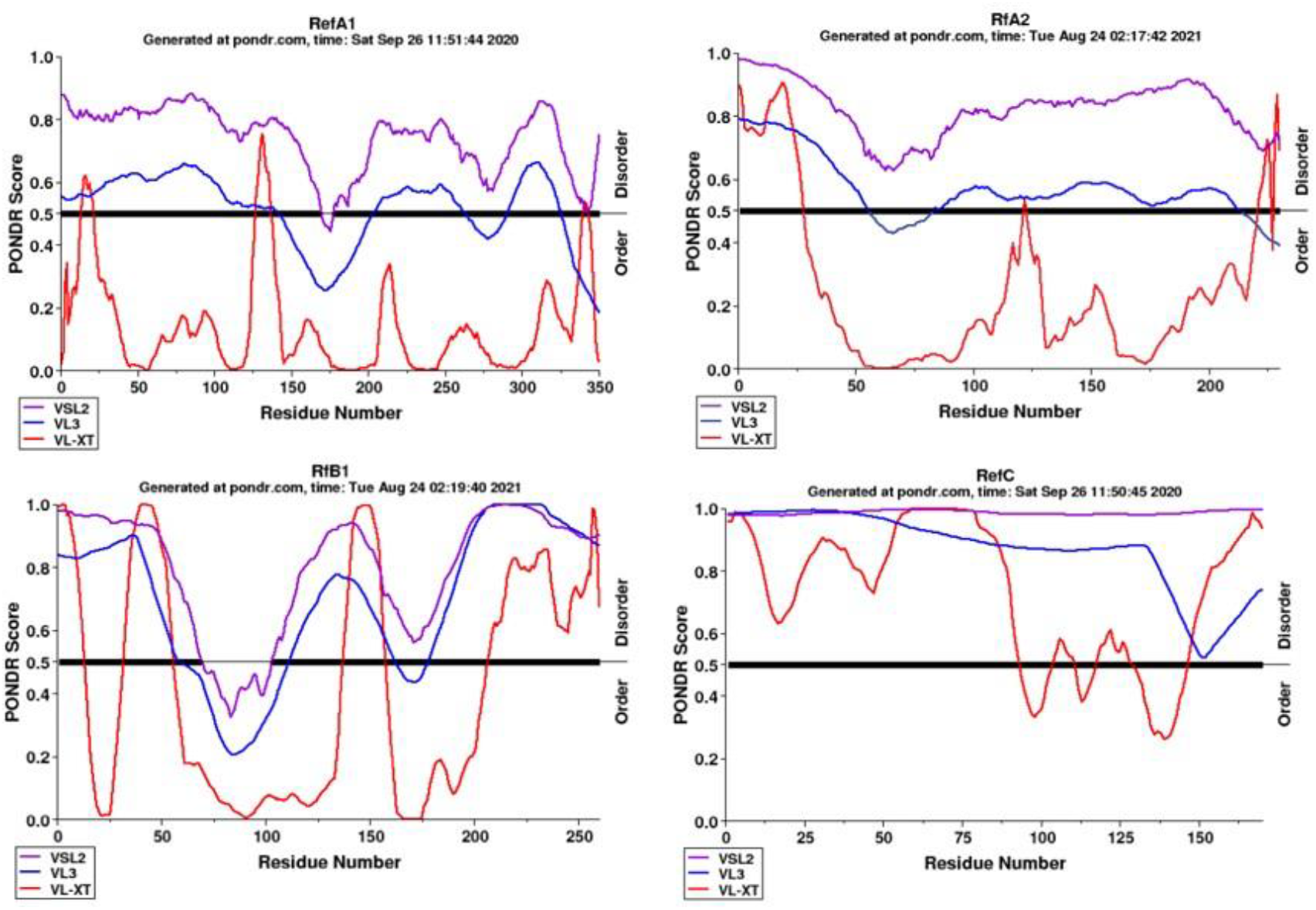
Disorder prediction of RfA1 by PONDER.

**Fig.2.**
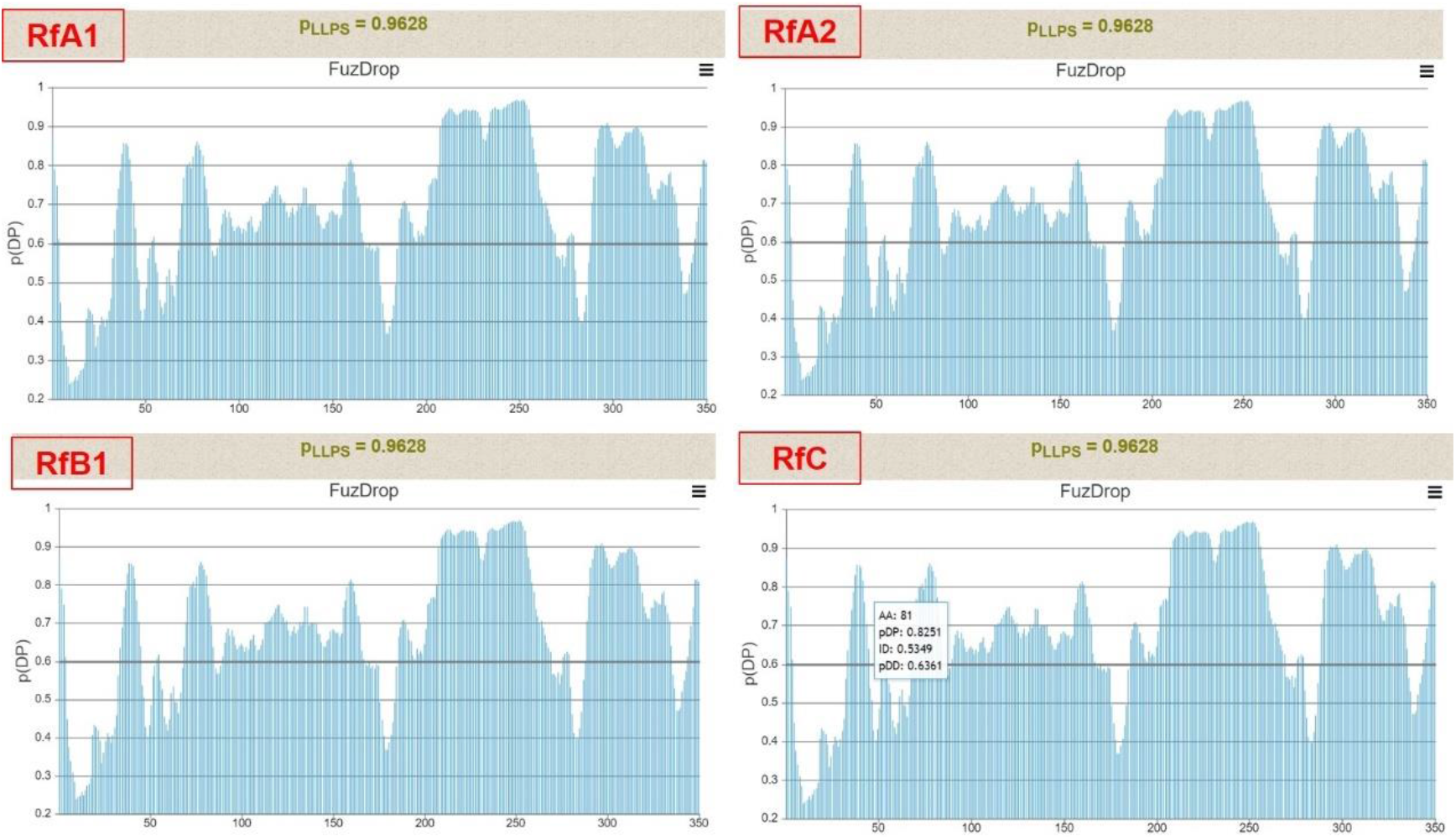
The probability of reflectins undergoes LLPS spontaneously, by FuzDrop.

**Fig.3.**
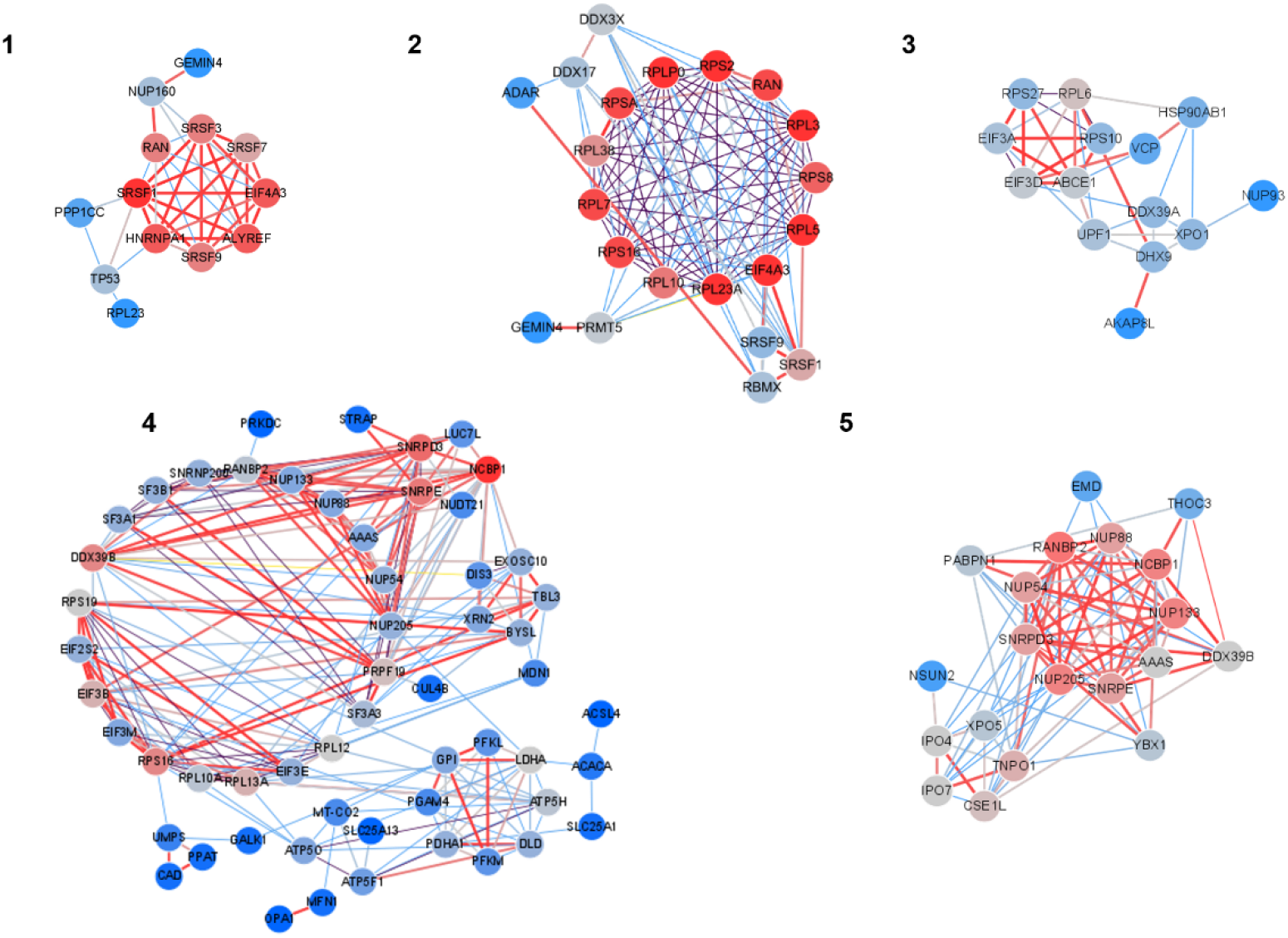
A protein-protein interaction network was constructed by the Metascape database. The network was visualized by Cytoscape 3.7.1 software. Built-in app MCODE was utilized to find clusters in the whole network. Five clusters were found and marked in the figure. Networks “1”, “2”,”3”, “4”, “5” correspond to Set1-1, Set1-2, Set5-3, Set3-4, Set3-5 in main text Fig. 5.

**Fig.4.**
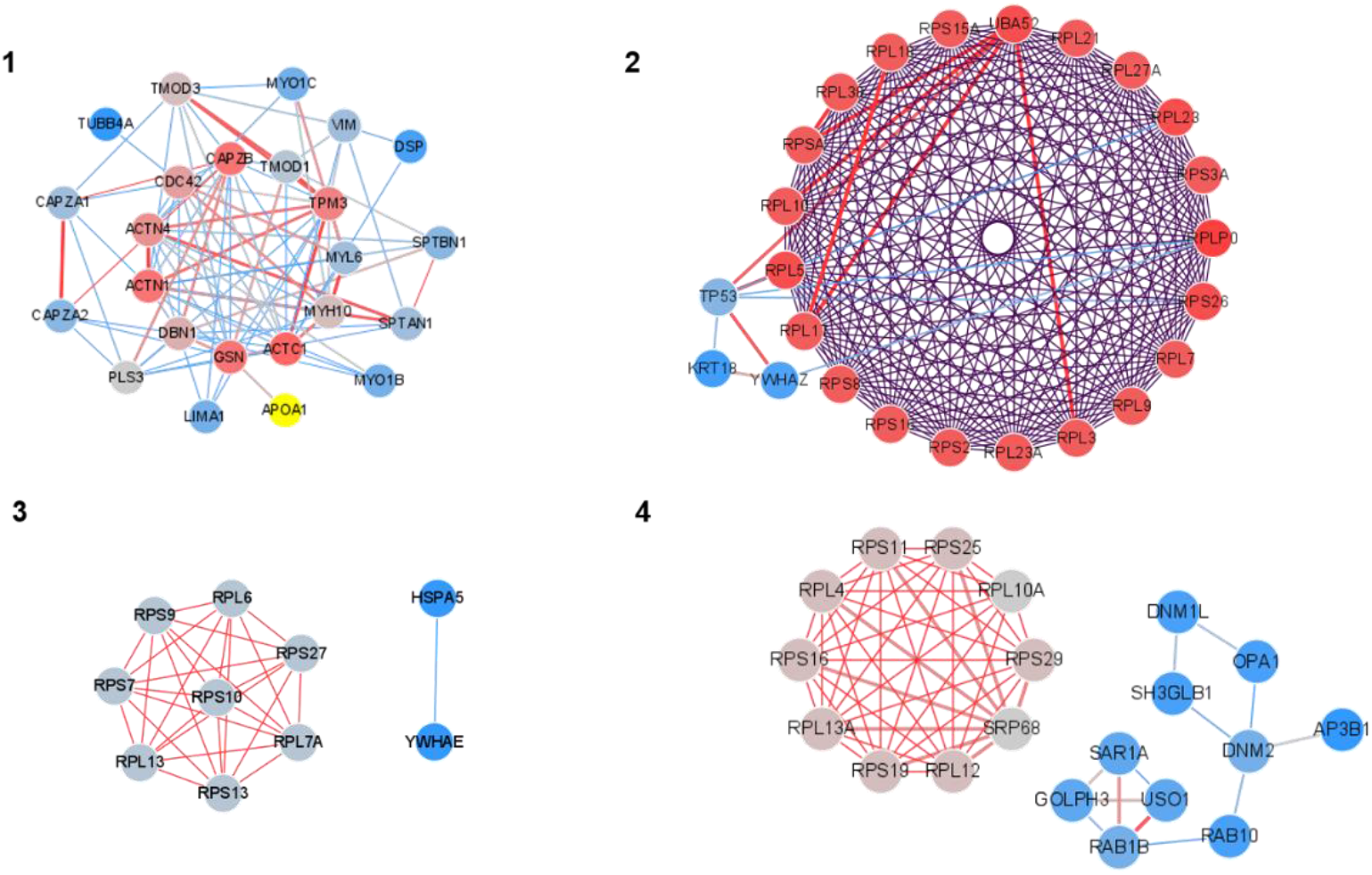
A protein-protein interaction network was constructed by the Metascape database. The network was visualized by Cytoscape 3.7.1 software. Built-in app MCODE was utilized to find clusters in the whole network. Five clusters were found and marked in the figure. Networks “1”, “2”,”3”, “4”, “5” correspond to Set1-1, Set1-2, Set5-3, Set3-4 in main text Fig. 6.

## References

[1] R.J.C.B. Hanlon, Cephalopod dynamic camouflage. Curr Biol 17 (2007) R400–R404.

[2] R. Hanlon, C.-C. Chiao, L. Mäthger, A. Barbosa, K. Buresch, and C.J.P.T.o.t.R.S.B.B.S. Chubb, Cephalopod dynamic camouflage: bridging the continuum between background matching and disruptive coloration. Philos T R Soc B 364 (2009) 429–437.

[3] A.R. Tao, D.G. DeMartini, M. Izumi, A.M. Sweeney, A.L. Holt, and D.E.J.B. Morse, The role of protein assembly in dynamically tunable bio-optical tissues. Biomaterials 31 (2010) 793–801.

[4] D.G. DeMartini, D.V. Krogstad, and D.E. Morse, Membrane invaginations facilitate reversible water flux driving tunable iridescence in a dynamic biophotonic system. Proc Natl Acad Sci U S A 110 (2013) 2552–6.

[5] T.L. Williams, S.L. Senft, J. Yeo, F.J. Martin-Martinez, A.M. Kuzirian, C.A. Martin, C.W. DiBona, C.T. Chen, S.R. Dinneen, H.T. Nguyen, C.M. Gomes, J.J.C. Rosenthal, M.D. MacManes, F. Chu, M.J. Buehler, R.T. Hanlon, and L.F. Deravi, Dynamic pigmentary and structural coloration within cephalopod chromatophore organs. Nat Commun 10 (2019) 1004.

[6] L.M. Mäthger, S.L. Senft, M. Gao, S. Karaveli, G.R. Bell, R. Zia, A.M. Kuzirian, P.B. Dennis, W.J. Crookes□Goodson, and R.R.J.A.F.M. Naik, Bright white scattering from protein spheres in color changing, flexible cuttlefish skin. ADV FUNCT MATER 23 (2013) 3980–3989.

[7] R.T. Hanlon, L.M. Mathger, G.R.R. Bell, A.M. Kuzirian, and S.L. Senft, White reflection from cuttlefish skin leucophores. Bioinspir Biomim 13 (2018) 035002.

[8] D.G. DeMartini, A. Ghoshal, E. Pandolfi, A.T. Weaver, M. Baum, and D.E.J.J.o.E.B. Morse, Dynamic biophotonics: female squid exhibit sexually dimorphic tunable leucophores and iridocytes. J Exp Biol 216 (2013) 3733–3741.

[9] D.G. DeMartini, M. Izumi, A.T. Weaver, E. Pandolfi, and D.E.J.J.o.B.C. Morse, Structures, organization, and function of reflectin proteins in dynamically tunable reflective cells. J Biol Chem 290 (2015) 15238–15249.

[10] R. Levenson, C. Bracken, C. Sharma, J. Santos, C. Arata, B. Malady, and D.E.J.J.o.B.C. Morse, Calibration between trigger and color: Neutralization of a genetically encoded coulombic switch and dynamic arrest precisely tune reflectin assembly. J Biol Chem 294 (2019) 16804–16815.

[11] Z. Guan, T. Cai, Z. Liu, Y. Dou, X. Hu, P. Zhang, X. Sun, H. Li, Y. Kuang, Q. Zhai, H. Ruan, X. Li, Z. Li, Q. Zhu, J. Mai, Q. Wang, L. Lai, J. Ji, H. Liu, B. Xia, T. Jiang, S.-J. Luo, H.-W. Wang, and C. Xie, Origin of the Reflectin Gene and Hierarchical Assembly of Its Protein. Curr Biol 27 (2017) 2833–2842.e6.

[12] R. Levenson, C. Bracken, N. Bush, and D.E. Morse, Cyclable Condensation and Hierarchical Assembly of Metastable Reflectin Proteins, the Drivers of Tunable Biophotonics. J Biol Chem 291 (2016) 4058–4068.

[13] L. Phan, D.D. Ordinario, E. Karshalev, W.G. Walkup IV, M.A. Shenk, and A.A.J.J.o.M.C.C. Gorodetsky, Infrared invisibility stickers inspired by cephalopods. J. MATER. CHEM. C 3 (2015) 6493–6498.

[14] P.B. Dennis, K.M. Singh, M.C. Vasudev, R.R. Naik, and W.J.J.A.M. Crookes-Goodson, Research update: a minimal region of squid reflectin for vapor-induced light scattering. APL MATER 5 (2017) 120701.

[15] G. Qin, P.B. Dennis, Y. Zhang, X. Hu, J.E. Bressner, Z. Sun, W.J. Crookes□Goodson, R.R. Naik, F.G. Omenetto, and D.L.J.J.o.P.S.P.B.P.P. Kaplan, Recombinant reflectin□based optical materials. J POLYM SCI POL PHYS 51 (2013) 254–264.

[16] D.D. Ordinario, L. Phan, W.G. Walkup IV, J.-M. Jocson, E. Karshalev, N. Hüsken, and A.A.J.N.c. Gorodetsky, Bulk protonic conductivity in a cephalopod structural protein. Nat Chem 6 (2014) 596–602.

[17] L. Phan, R. Kautz, E.M. Leung, K.L. Naughton, Y. Van Dyke, and A.A.J.C.o.M. Gorodetsky, Dynamic materials inspired by cephalopods. Chem Mater 28 (2016) 6804–6816.

[18] C. Yu, Y. Li, X. Zhang, X. Huang, V. Malyarchuk, S. Wang, Y. Shi, L. Gao, Y. Su, and Y.J.P.o.t.N.A.o.S. Zhang, Adaptive optoelectronic camouflage systems with designs inspired by cephalopod skins. Proc Natl Acad Sci U S A 111 (2014) 12998–13003.

[19] J. Song, R. Levenson, J. Santos, L. Velazquez, F. Zhang, D. Fygenson, W. Wu, and D.E.J.L. Morse, Reflectin Proteins Bind and Reorganize Synthetic Phospholipid Vesicles. Langumir 36 (2020) 2673–2682.

[20] A. Chatterjee, J.A. Cerna Sanchez, T. Yamauchi, V. Taupin, J. Couvrette, and A.A. Gorodetsky, Cephalopod-inspired optical engineering of human cells. Nat Commun 11 (2020) 2708.

[21] J. Ogawa, Y. Iwata, N.U. Tonnu, C. Gopinath, L. Huang, S. Itoh, R. Ando, A. Miyawaki, I.M. Verma, and G.M.J.b. Pao, Genetic manipulation of the optical refractive index in living cells. bioRxiv (2020).

[22] J.F. Dishinger, H.L. Kee, P.M. Jenkins, S. Fan, T.W. Hurd, J.W. Hammond, Y.N.-T. Truong, B. Margolis, J.R. Martens, and K.J.J.N.c.b. Verhey, Ciliary entry of the kinesin-2 motor KIF17 is regulated by importin-ß2 and RanGTP. Nature Cell Biology 12 (2010) 703–710.

[23] H. Ishikawa, and W.F.J.C.S.H.p.i.b. Marshall, Intraflagellar transport and ciliary dynamics. Cold Spring Harbor perspectives in biology 9 (2017) a021998.

[24] H.L. Kee, J.F. Dishinger, T.L. Blasius, C.-J. Liu, B. Margolis, and K.J.J.N.c.b. Verhey, A size-exclusion permeability barrier and nucleoporins characterize a ciliary pore complex that regulates transport into cilia. Nature Cell Biology 14 (2012) 431–437.

[25] D.K. Breslow, E.F. Koslover, F. Seydel, A.J. Spakowitz, and M.V.J.J.o.C.B. Nachury, An in vitro assay for entry into cilia reveals unique properties of the soluble diffusion barrier. Journal of Cell Biology 203 (2013) 129–147.

[26] O.D. Weiner, W.A. Marganski, L.F. Wu, S.J. Altschuler, and M.W. Kirschner, An actin-based wave generator organizes cell motility. PLoS Biol 5 (2007) e221.

[27] E.S. Chhabra, and H.N.J.N.c.b. Higgs, The many faces of actin: matching assembly factors with cellular structures. Nature cell biology 9 (2007) 1110–1121.

[28] M.R. Mejillano, S. Kojima, D.A. Applewhite, F.B. Gertler, T.M. Svitkina, and G.G. Borisy, Lamellipodial versus filopodial mode of the actin nanomachinery: pivotal role of the filament barbed end. Cell 118 (2004) 363–73.

[29] M. Innocenti, New insights into the formation and the function of lamellipodia and ruffles in mesenchymal cell migration. Cell Adh Migr 12 (2018) 401–416.

[30] W.J. Crookes, L.-L. Ding, Q.L. Huang, J.R. Kimbell, J. Horwitz, and M.J.J.S. McFall-Ngai, Reflectins: the unusual proteins of squid reflective tissues. Science 303 (2004) 235–238.

[31] B. Xue, R.L. Dunbrack, R.W. Williams, A.K. Dunker, V.N.J.B.e.B.A.-P. Uversky, and Proteomics, PONDR-FIT: a meta-predictor of intrinsically disordered amino acids. BBA-PROTEINS PROTEOM 1804 (2010) 996–1010.

[32] L.J.J.B. McGuffin, Intrinsic disorder prediction from the analysis of multiple protein fold recognition models. Bioinformatics 24 (2008) 1798–1804.

[33] J.J. Ward, J.S. Sodhi, L.J. McGuffin, B.F. Buxton, and D.T.J.J.o.m.b. Jones, Prediction and functional analysis of native disorder in proteins from the three kingdoms of life. J MOL BIOL 337 (2004) 635–645.

[34] I. Walsh, A.J. Martin, T. Di Domenico, and S.C.J.B. Tosatto, ESpritz: accurate and fast prediction of protein disorder. Bioinformatics 28 (2012) 503–509.

[35] M. Hardenberg, A. Horvath, V. Ambrus, M. Fuxreiter, and M.J.P.o.t.N.A.o.S. Vendruscolo, Widespread occurrence of the droplet state of proteins in the human proteome. Proc Natl Acad Sci U S A 117 (2020) 33254–33262.

[36] R. Roy, D. Durie, H. Li, B.-Q. Liu, J.M. Skehel, F. Mauri, L.V. Cuorvo, M. Barbareschi, L. Guo, and M.J.N.a.r. Holcik, hnRNPA1 couples nuclear export and translation of specific mRNAs downstream of FGF-2/S6K2 signalling. NUCLEIC ACIDS RES 42 (2014) 12483–12497.

[37] W.M. Michael, M. Choi, and G.J.C. Dreyfuss, A nuclear export signal in hnRNP A1: a signal-mediated, temperature-dependent nuclear protein export pathway. Cell 83 (1995) 415–422.

[38] G.M. Hautbergue, L.M. Castelli, L. Ferraiuolo, A. Sanchez-Martinez, J. Cooper-Knock, A. Higginbottom, Y.-H. Lin, C.S. Bauer, J.E. Dodd, and M.A.J.N.c. Myszczynska, SRSF1 -dependent nuclear export inhibition of C9ORF72 repeat transcripts prevents neurodegeneration and associated motor deficits. Nat Commun 8 (2017) 1–18.

[39] T. Güttler, and D.J.T.E.j. Görlich, Ran dependent nuclear export mediators: a structural perspective. EMBO J 30 (2011) 3457–3474.

[40] M.K. Connor, R. Kotchetkov, S. Cariou, A. Resch, R. Lupetti, R.G. Beniston, F. Melchior, L. Hengst, and J.M.J.M.b.o.t.c. Slingerland, CRM1/Ran-mediated nuclear export of p27Kip1 involves a nuclear export signal and links p27 export and proteolysis. MOL BIOL CELL 14 (2003) 201–213.

[41] R. Zhang, R. Mehla, and A.J.P.o. Chauhan, Perturbation of host nuclear membrane component RanBP2 impairs the nuclear import of human immunodeficiency virus-1 preintegration complex (DNA). PLOS ONE 5 (2010) e15620.

[42] B. Cautain, R. Hill, N. de Pedro, and W.J.T.F.j. Link, Components and regulation of nuclear transport processes. FEBS J 282 (2015) 445–462.

[43] A.P. Sfakianos, A.J. Whitmarsh, and M.P.J.B.S.T. Ashe, Ribonucleoprotein bodies are phased in. Biochem Soc Trans 44 (2016) 1411–1416.

[44] N. Mukherjee, H.-H. Wessels, S. Lebedeva, M. Sajek, M. Ghanbari, A. Garzia, A. Munteanu, D. Yusuf, T. Farazi, and J.I.J.N.a.r. Hoell, Deciphering human ribonucleoprotein regulatory networks. NUCLEIC ACIDS RES 47 (2019) 570–581.

[45] T.D. Pollard, Actin and Actin-Binding Proteins. Cold Spring Harb Perspect Biol 8 (2016).

[46] R. Uribe, and D. Jay, A review of actin binding proteins: new perspectives. Mol Biol Rep 36 (2009) 121–5.

[47] R. Dominguez, and K.C. Holmes, Actin structure and function. Annu Rev Biophys 40 (2011) 169–86.

[48] P.W. Gunning, U. Ghoshdastider, S. Whitaker, D. Popp, and R.C.J.J.o.c.s. Robinson, The evolution of compositionally and functionally distinct actin filaments. J Cell Sci 128 (2015) 2009–2019.

[49] Y. Zhou, B. Zhou, L. Pache, M. Chang, A.H. Khodabakhshi, O. Tanaseichuk, C. Benner, and S.K.J.N.c. Chanda, Metascape provides a biologist-oriented resource for the analysis of systems-level datasets. NAT METHODS 10 (2019) 1–10.

[50] E.A. Foley, and T.M.J.N.r.M.c.b. Kapoor, Microtubule attachment and spindle assembly checkpoint signalling at the kinetochore. NAT REV MOL CELL BIO 14 (2013) 25–37.

[51] T. Wittmann, A. Hyman, and A.J.N.c.b. Desai, The spindle: a dynamic assembly of microtubules and motors. NAT CELL BIOL 3 (2001) E28–E34.

[52] L.G. Glotfelty, A. Zahs, C. Iancu, L. Shen, and G.A.J.A.J.o.P.-C.P. Hecht, Microtubules are required for efficient epithelial tight junction homeostasis and restoration. American Journal of Physiology-Cell Physiology 307 (2014) C245–C254.

[53] K. Sugioka, and H.J.C.o.i.c.b. Sawa, Formation and functions of asymmetric microtubule organization in polarized cells. Current opinion in cell biology 24 (2012) 517–525.

[54] R. Levenson, D.G. DeMartini, and D.E.J.A.M. Morse, Molecular mechanism of reflectin’s tunable biophotonic control: Opportunities and limitations for new optoelectronics. APL Mater 5 (2017) 104801.

[55] J. Schindelin, I. Arganda-Carreras, E. Frise, V. Kaynig, M. Longair, T. Pietzsch, S. Preibisch, C. Rueden, S. Saalfeld, and B.J.N.m. Schmid, Fiji: an open-source platform for biological-image analysis. Nat Methods 9 (2012) 676–682.

[56] G. Yu, L.-G. Wang, Y. Han, and Q.-Y.J.O.a.j.o.i.b. He, clusterProfiler: an R package for comparing biological themes among gene clusters. OMICS 16 (2012) 284–287.

[57] B.J. Perez-Riverol Y, Bandla C, Hewapathirana S, García-Seisdedos D, Kamatchinathan S, Kundu D, Prakash A, Frericks-Zipper A, Eisenacher M, Walzer M, Wang S, Brazma A, Vizcaíno JA The PRIDE database resources in 2022: A Hub for mass spectrometry-based proteomics evidences. Nucleic Acids Res 50(D1):D543–D552 (2022).

